# Haplotype-resolved genome assembly enables gene discovery in the red palm weevil *Rhynchophorus ferrugineus*

**DOI:** 10.1101/2020.08.30.273383

**Authors:** Guilherme B. Dias, Musaad A. Altammami, Hamadttu A.F. El-Shafie, Fahad M. Alhoshani, Mohamed B. Al-Fageeh, Casey M. Bergman, Manee M. Manee

## Abstract

The red palm weevil *Rhynchophorus ferrugineus* (Coleoptera: Curculionidae) is an economically-important invasive species that attacks multiple species of palm trees around the world. A better understanding of gene content and function in *R. ferrugineus* has the potential to inform pest control strategies and thereby mitigate economic and biodiversity losses caused by this species. Using 10x Genomics linked-read sequencing, we produced a haplotype-resolved diploid genome assembly for *R. ferrugineus* from a single heterozygous individual with modest sequencing coverage (~62x). Benchmarking against conserved single-copy Arthropod orthologs suggests both pseudo-haplotypes in our *R. ferrugineus* genome assembly are highly complete with respect to gene content, and do not suffer from haplotype-induced duplication artifacts present in a recently published hybrid assembly for this species. Annotation of the larger pseudo-haplotype in our assembly provides evidence for 23,413 protein-coding loci in *R. ferrugineus*, including over 13,000 predicted proteins annotated with Gene Ontology terms and over 6,000 loci independently supported by high-quality Iso-Seq transcriptomic data. Our assembly also includes 95% of *R. ferrugineus* chemosensory, detoxification and neuropeptide-related transcripts identified previously using RNA-seq transcriptomic data, and provides a platform for the molecular analysis of these and other functionally-relevant genes that can help guide management of this widespread insect pest.

## Introduction

The insect order Coleoptera (beetles) is one of the most diverse groups of organisms on earth, with over 350,000 species currently described and an estimated 1.5 million species in total^1^. Despite this unrivaled organismal diversity, there has not yet been an inordinate fondness for studying beetle genomes^2^, with only 37 species having genome assemblies in NCBI as of August 2020. The family Curculionidae (“true” weevils) is one of the largest beetle groups, containing over 80,000 described species including many important agricultural pests. Among these, the red palm weevil (RPW) *Rhynchophorus ferrugineus* is a widespread invasive species that attacks a variety of palm tree species. The RPW is of particular interest since it is the major arthropod pest of the date palm (*Phoenix dactylifera*), resulting in economic losses in the order of tens of millions of dollars annually^3^.

RPW adult females bore into palm trees to deposit their eggs, wherein larvae hatch and consume the surrounding trunk tissue causing extensive damage as they develop. This trait of being a “concealed tissue borer” makes infestation hard to detect in its early stages and often results in the death of infected plants. Concealed boring also protects developing larvae from abiotic stressors, and facilitates weevil dispersion across large distances during commercialization of palm offshoots for farming and ornamental purposes^4^. These factors, together with polyphagy and strong flight ability^5^, contribute to the invasive potential and economic impact of the RPW.

To reduce the economic and biodiversity losses caused by the RPW, there has been growing interest to identify RPW genes that can be used to guide strategies for pest management in this species. Previous gene discovery efforts for the RPW have mainly relied on transcriptome data, using different sequencing platforms, as well as a range of tissues, developmental stages, and strains^6–11^. Transcriptomics is a cost-effective strategy of gene discovery compared to whole-genome sequencing since only a fraction of the genome is represented in mature transcripts. However, the time- and tissue-specificity of gene expression makes it hard to capture all protein-coding genes in an organism using a limited number of RNA-seq samples. In addition, aspects of transcript structure and gene organization cannot be inferred from transcriptome assemblies alone.

Like many agriculturally important non-model species, efforts to generate a genome assembly for the RPW to aid gene discovery have been hampered by the heterozygosity inherent in diploid organisms. However, advances in genomics now allow resolution of both haplotypes in *de novo* assemblies of diploid organisms, typically using either linked (e.g. 10x Genomics) or long (e.g. PacBio or Oxford Nanopore) reads^12^. Recently, a hybrid assembly using a combination of Illumina, 10x Genomics and Oxford Nanopore sequencing was reported for the RPW that was used for gene discovery and analysis of gene family evolution^13^. This hybrid assembly reported an unusually high rate of gene family expansion in the RPW genome relative to other beetle species, and also reported a very high number of duplicated genes in the BUSCO gene set, which are expected to be present in a single copy in most organisms^14^. To overcome limitations of previous transcriptome-based gene discovery efforts in the RPW^6–11^ and to evaluate the correctness of the previously-reported RPW genome assembly^13^, here we report a haplotype-resolved (“phased”) diploid genome assembly from an independent RPW sample generated using 10x Genomics linked-read sequencing. We provide evidence that the previously-reported RPW genome hybrid assembly contains a large proportion of artifactually duplicated sequences that have arisen from multiple haplotypes being scaffolded into a single haploid representation of the genome^15^. We demonstrate that our haplotype-resolved diploid assembly does not suffer from such artifacts and therefore provides a more accurate resource for understanding the genome and gene content of this important agricultural pest.

## Materials and Methods

### Sample, library preparation and sequencing

A single three-week-old RPW larvae was selected randomly for sequencing from a colony of RPW reared on date palms of the ‘Khalas’ cultivar in the shade house at the Date Palm Research Center of Excellence at King Faisal University. This colony was established from multiple individuals sampled in February 2017 using insecticide-free pheromone traps in the Al-Ahsa oasis in Saudi Arabia.

The individual larvae selected for sequencing was sectioned into 4-8 mg pieces, one of which was used for DNA extraction following the 10x Genomics recommended protocol for single insect DNA purification (https://support.10xgenomics.com/permalink/7HBJeZucc80CwkMAmA4oQ2). This protocol uses a salting out method adapted from Miller *et al.*^16^. We chose larval tissue for sequencing since the recommended 10x Genomics DNA extraction protocol for insects yielded longer molecules for larval relative to adult tissues. As a consequence, the sex of the individual sequenced here was initially unknown but was later determined to most likely be female (see Results and Discussion).

Purified genomic DNA was size selected to remove fragments shorter than 20 kb using the BluePippin instrument (Sage Science). After size selection, 0.6 ng of DNA was loaded onto the 10x Genomics Chromium Genome Chip for gel bead-in-emulsion generation, barcoding, and library construction using the 10x Genomics Chromium Genome Reagent Kit Protocol v2 (RevB). DNA sequencing was carried out on a Illumina NextSeq 500 mid-output flow cell with 150 bp paired-end (PE) read layout.

### Genome assembly

Raw sequencing reads were assembled using the Supernova assembler v2.1.1 with default parameters and exported to fasta format using the ‘pseudohap2’ style^17^. This output mode generates two pseudo-haplotype assemblies that differ in genomic regions where maternal and paternal haplotypes can be phased (“phase blocks”) but are identical in homozygous and unphased blocks of the genome (Supplementary Figure S1). Pseudo-haplotype1 was selected as the main assembly for analysis and annotation since it is slightly longer than pseudo-haplotype2. Both pseudo-haplotypes were deposited in GenBank (see Data availability section for accession numbers). To better understand differences between our pseudo-haplotype assemblies and the hybrid assemblies of Hazzouri *et al.*^13^, we also exported our RPW Supernova assembly in ‘megabubbles’ style^17^ which includes maternal and paternal phase blocks together with unphased blocks in a single file (Supplementary Figure S1).

Contigs in 11 scaffolds from both pseudo-haplotypes were trimmed to remove small (<50 bp) internal adapter sequences flanking assembly gaps that were identified in NCBI’s contamination screen report. Redundancy in the Supernova assembly was eliminated using the ‘sequniq’ command from GenomeTools v1.5.9^18^. These filtering steps resulted in the removal of two contaminated and 3,694 redundant scaffolds spanning 10,665,716 bp, or 1.78% of the original Supernova assembly size.

### Genome annotation

Prior to genome annotation, a custom repeat library was generated from pseudo-haplotype1 using RepeatModeler v1.0.11 (-engine ncbi) and used to soft mask the pseudo-haplotype1 assembly with RepeatMasker v4.0.9 (-gff -u -a -s -no_is -xsmall -e ncbi) (http://www.repeatmasker.org/). Protein-coding gene annotation of pseudo-haplotype1 was performed with BRAKER v2.1.5^19^, an automated gene annotation pipeline that uses extrinsic evidence in the form of spliced alignments from RNA-seq data and protein sequences to train and predict gene structures with AUGUSTUS^20–22^.

Short-read RNA-seq data from *R. ferrugineus* for training BRAKER was obtained from the NCBI SRA database. Three different datasets were used consisting of Illumina short-read sequencing of: (i) polyA+ RNA from pooled male and female adults (PRJDB3020)^7^, (ii) total RNA from pooled male and female antennae (PRJNA275430)^9^, and (iii) total RNA from RPW larvae, pupae and adults of both sexes (PRJNA598560)^10^. Raw RNA-seq reads were quality filtered with fastp v0.20.0^23^ using default parameters, and aligned to pseudo-haplotype1 using HiSat2 v2.1.0 (--dta)^24^. Resulting alignments were position sorted and converted to BAM format using SAMtools v1.9^25^. Spliced alignments of predicted protein sequences from 15 Coleopteran species plus *Drosophila melanogaster* to the RPW pseudo-haplotype1 assembly were obtained with ProtHint v2.4.0 pipeline^26^, and were also used to train BRAKER. Species names and NCBI accession numbers of genomes used as sources of predicted protein sequences for BRAKER training are listed in Supplementary Table S1. Genome-wide protein coding gene annotation was performed using BRAKER v2.1.5 (--prg=ph --etpmode --softmasking)^19^ and the gene set was based on AUGUSTUS predictions only. Five genes with internal stop codons were then removed from the BRAKER AUGUSTUS predictions to generate the final annotation submitted to NCBI.

Protein-coding gene models from BRAKER output were functionally annotated through protein signature scanning and sequence similarity searches against multiple databases. InterProScan v5.32-71.0^27^ was used to search the InterPro v71.0 member databases, and Diamond v0.9.32^28^ was used to search the non-redundant (*nr*) protein database from NCBI (from June 2020). The resulting similarity hits from InterPro and *nr* were imported to Blast2GO v5.2.5^29^ for final annotation with Gene Ontology (GO) terms^30^. Blast2GO was used to: (i) retrieve GO terms associated with *nr* protein similarity hits (mapping pipeline), (ii) annotate sequences with the most specific and reliable GO terms available from the mapping step, (iii) merge InterProScan associated GO IDs to the annotation, and (iv) augment the final annotation with the newly incorporated InterProScan GO IDs. All Blast2GO pipelines were run with default settings.

### Evaluation of assembly quality

General metrics for unmasked versions of both pseudo-haplotypes of our phased genome assembly, the megabubbles version of our phased genome assembly, assemblies from Hazzouri *et al.*^13^ (David Nelson, personal communication; GCA_012979105.1) and the *Tribolium castaneum* reference genome (GCF_000002335.3)^31^ were collected with the ‘stats.sh’ utility script from BBMap v38.76^32^. Completeness of unmasked genome assemblies was assessed with BUSCO v4.0.6 (-m genome -l arthropoda_-odb10 --augustus_species tribolium2012)^14^ using the Arthropoda gene set from OrthoDB v10^33^. Nucleotide differences between pseudo-haplotypes in our RPW assembly were computed by aligning orthologous scaffolds with minimap2 v2.17 (-cx asm20 --cs --secondary=no)^34^ and extracting variants with paftools.js call across alignments at different minimal length cutoffs (-L) of 1 kb, 10 kb, and 50 kb. The total length of phased blocks from each pseudo-haplotype was calculated from the Supernova index files. To visualize heterozygosity along phase blocks we aligned the raw 10x data produced here to pseudo-haplotype1 with BWA-MEM v0.7.17-r118, removed alignments with MAPQ=0 using SAMtools v1.9^25^, called variants with BCFtools v1.9^25^ (call -v -m) and VCFtools v0.1.16^35^ (--remove-indels --remove-filtered-all --recode --recode-INFO-all), and calculated the B-allele frequency of variants using the information in the DP4 field from the resulting VCF file. Single-nucleotide variants and phase blocks were visualized for the 10 longest scaffolds using karyoploteR v1.10.2^36^. To identify potential sex chromosome scaffolds and determine the sex of the individual sequenced, we subsampled male and female Illumina reads from Hazzouri *et al.*^13^ (SRX5416728, SRX5416729) and the 10x Genomics reads produced here (SRX7520800) to ~39 Gb using seqtk v1.3 (https://github.com/lh3/seqtk), aligned to pseudo-haplotype1 using BWA-MEM v0.7.17-r1188^37^, removed alignments contained within repeat-masked regions or with MAPQ=0 using SAMtools v1.9^25^, calculated the mapped read depth using BEDtools v2.29.0^38^ (genomecov -dz), and finally calculated the ratio of male/female mean mapped read depth for each scaffold. The mean mapped read depth across the 10 longest scaffolds in pseudo-haplotype1 was visualized with karyoploteR v1.10.2^36^. Estimates of total genome size from unassembled Illumina reads were generated using findGSE v1.94^39^ and GenomeScope v1.0.0^40^. Frequency histograms for 21-mers were obtained with Jellyfish v2.3.0 with a max k-mer coverage of 1,000,000^41^.

To test whether unassembled Illumina DNA-seq datasets supported the high proportion of BUSCO genes that are classified as duplicated in the final Hazzouri *et al.*^13^ hybrid assembly (M_pseudochr; GCA_012979105.1), we first classified arthropod BUSCO genes as being single-copy or duplicated in the M_pseudochr assembly using BUSCO v4.0.6 (-m genome -l arthropoda_odb10 --augustus_species tribolium2012)^14^. Next, we aligned unassembled DNA-seq reads generated in this study (SRX7520800) and in Hazzouri *et al.*^13^ (SRX5416727, SRX5416728, SRX5416729) to our unmasked pseudo-haplotype1 assembly using BWA-MEM v0.7.17-r1188 with default parameters^37^. The resulting alignments were position sorted and low quality mappings (MAPQ = 0) were removed using SAMtools v1.9^25^. Mean depth of all mapped reads or only high-quality mapped reads were then calculated using BEDtools v2.29.0 (bedtools coverage -mean)^38^ across the complete genomic range of each BUSCO gene in our pseudo-haplotype1 assembly. Finally, histograms of mean read depth for BUSCO genes in our pseudo-haplotype1 assembly (partitioned as being single-copy or duplicated based on classification in the M_pseudochr assembly were plotted for each DNA-seq dataset using R and ggplot2^42, 43^. The cumulative distribution functions for mean mapped read depth were compared between single-copy and duplicated gene sets for each DNA-seq dataset using the Kolmogorov–Smirnov test in R v3.6.3 with the alternative hypothesis that the mean mapped read depth of duplicated BUSCO genes is greater than the mean mapped read depth of single-copy BUSCO genes (*α* = 0.05).

### Evaluation of annotation quality

Support for exon-intron junctions in gene models from our BRAKER annotation was evaluated against the spliced RNA-seq alignments using paftools.js from minimap2 v2.17^34^. Completeness of the following RPW and *T. castaneum* gene sets was evaluated using BUSCO v4.0.6 (-m transcriptome -l arthropoda_odb10 --augustus_species tribolium2012)^14^: BRAKER annotation of our pseudo-haplotype1 assembly, a re-processed version of the long read transcriptome dataset from Yang *et al.*^10^ (see below), the Funannotate annotation of the M_v.1 hybrid assembly from Hazzouri *et al.*^13^ (Khaled Hazzouri, personal communication), BRAKER re-annotation of the M_pseudochr from Hazzouri *et al.*^13^ using the same procedure used for our pseudo-haplotype1 (see above), and the *T. castaneum* v5.2 reference genome annotation from Shelton *et al.*^31^. Two BUSCO analyses were performed for each gene set, one using all transcripts and one using a single randomly chosen isoform per locus after clustering BRAKER transcripts into distinct loci using GffRead 0.11.7 (--cluster-only)^44^. Clustering transcripts with GffRead takes into account strand orientation and exon-intron structure, and although this process might group transcripts that only partially overlap it its more specific than a simple interval overlap. The “all transcripts” BUSCO analysis provides an upper bound estimate of BUSCO recovery in a gene set, since one isoform of a locus might fit a BUSCO gene profile better than another isoform, but can falsely classify BUSCO genes as being duplicated due to the identification of multiple isoforms of the same locus. The “one isoform per locus” BUSCO analysis provides a conservative estimate of BUSCO gene recovery which more accurately represents the true degree of duplication in the gene set that is not biased by alternative isoform usage.

Our BRAKER genome annotation was also evaluated against two external datasets of RPW genes. First, we used a dataset of full-length cDNA transcripts reported in Yang *et al.*^10^ generated using PacBio long-read isoform sequencing (Iso-Seq) from mixed tissues (larvae, pupae, and adults of both sexes). We observed anti-sense artifacts in the processed Iso-Seq transcriptomes reported by Yang *et al.*^10^ (High quality: SRX7519788; Low quality: SRX8694670) (Supplementary Figure S2). Thus we re-processed the original circular consensus sequences (CCS) from Yang *et al.*^10^ (SRX7495110) using default parameters of the isoseq3 pipeline in SMRT Link v8.0.0.79519. Unpolished isoform consensus sequences output from isoseq3 cluster were polished with Illumina RNA-seq reads from Yang *et al.*10 using Lordec v0.9^45^ based on best performing parameters from Hu *et al.*^46^ (-k 21 -t 15 -b 1000 -e 0.45 -s 5). Polished isoform consensus sequences were then aligned to our pseudo-haplotype1 assembly using minimap2 v2.17 (-ax splice --cs --secondary=no)^34^. Supplementary alignments were then removed using SAMtools v1.9 to retain only primary and representative alignments. The resulting spliced alignments were position sorted and converted to GTF2 format using SAMtools v1.9, BEDtools v2.29.0, and UCSC tools v377^25, 38, 47^. Finally, we clustered re-processed Iso-Seq transcripts into distinct loci using GffRead v0.11.7 (--cluster-only), then compared Iso-Seq loci with BRAKER loci using GffCompare v0.11.2^44^. GffCompare identifies multiple types of overlaps between a reference set of transcripts and a query. These include overlaps representing the perfect concordance of the exon-intron structure, but also partial intronic and exonic overlaps as well as the containment of query transcripts by reference transcripts and vice-versa. Exonic overlaps between reference and query transcripts in opposite strands are flagged but do not contribute to the final statistics of overlapped loci^44^.

Second, we compared our BRAKER genome annotation against curated sets of RPW genes that are potentially relevant for pest mitigation strategies^9, 11, 48^. Transcript identifiers for chemosensory genes were obtained from Antony *et al.*^9^ and parsed from their transcriptome assembly (GDKA01000000). Transcripts for cytochrome P450 monooxygenases were obtained from Antony *et al.*^48^. Transcripts for neuropeptides and their G-protein coupled receptors (GPCRs) were obtained from Zhang *et al.*^11^ (MK751489–MK751534, MK751535–MK751576). All transcripts for curated gene sets were aligned to our pseudo-haplotype1 assembly, converted to GTF2, and clustered into distinct loci as for the Iso-Seq transcripts above. Transcripts for curated genes that could not be mapped to the RPW pseudo-haplotype1 assembly were further analyzed by directly querying DNA-seq data generated in this study (SRX7520800) and in Hazzouri *et al.*^13^ (SRX5416727, SRX5416728, SRX5416729). DNA-seq reads were mapped directly to the transcript contigs using minimap2 v2.17 (-ax sr)^34^, sorted and converted to BAM format using SAMtools v1.9^25^. Mean mapped read depth over each entire transcript was then calculated using BEDtools v2.29.0 (coverage -mean)^38^. To correct strand orientation errors observed in the mapped RPW curated gene sets from Antony *et al.*^9^ (Supplementary Figure S2), we used the combined RNA-seq dataset (used above as input to BRAKER) to assemble a reference-guided transcriptome using StringTie v2.0^49^. We removed genes for which no strand could be computed by StringTie (mostly single-exon genes), then overlapped the location of mapped transcripts from Antony *et al.*^9^, Antony *et al.*^48^ and Zhang *et al.*^11^ with our StringTie transcripts using GffCompare v0.11.2 to obtain the corresponding StringTie transcript for each curated gene in order to evaluate their consistency with BRAKER gene models. Finally, we compared StringTie transcripts for curated genes with BRAKER loci using GffCompare v0.11.2^44^. We note that while both StringTie transcripts and BRAKER annotations use the same underlying mapped RNA-seq data as input, StringTie transcripts were not used as evidence in BRAKER training nor were BRAKER gene models used in StringTie assembly, and thus BRAKER and StringTie annotations represent independent predictions of transcript structure.

## Results and Discussion

### Haplotype-resolved diploid assembly using 10x Genomics linked reads provides an accurate representation of RPW genome content

We prepared a 10x Genomics linked-read sequencing library from a single RPW individual originating from Al-Ahsa, Saudi Arabia and used this library to generate over 145 million 150-bp PE Illumina reads, totaling ~40.4 Gb after adapter trimming. Using this data, we assembled a draft phased diploid genome assembly for *R. ferrugineus* using Supernova^17^. We exported our diploid assembly in ‘pseudohap2’ format (Supplementary Figure S1), which produces two output files each having a phased ‘pseudo-haplotype’ assembly. In regions where haplotype phasing can be achieved, maternal and paternal phase blocks are randomly assigned to one of the two pseudo-haplotype assemblies. In regions where phasing cannot be achieved, either because low heterozygosity or insufficient linked-read data, the two pseudo-haplotypes are identical.

Assembly statistics and BUSCO scores for both pseudo-haplotypes in our assembly are presented in Table 1. The total length of each pseudo-haplotype is approximately 590 Mb, with contig N50’s of nearly 38 kb, and scaffold N50’s of over 470 kb. Approximately 98% of Arthropod BUSCOs are found completely represented in both pseudo-haplotypes, ~96% of which are single copy and only ~2% are duplicated. The completeness of our RPW pseudo-haplotype assemblies is comparable to the current reference genome of the best studied beetle species *T. castaneum*^31^, which has 99.1% complete BUSCOs with 98.6% being single-copy. Over 140 Mb (~24%) of each pseudo-haplotype is phased (Supplementary Files 1 and 2), with the two pseudo-haplotypes differing by <0.4% at aligned orthologous sites, and the majority of differences being single nucleotide polymorphisms and short indels (Supplementary Table S2). Because the two pseudo-haplotypes produced in our assembly are very similar, we chose pseudo-haplotype1 – the longer of the two assemblies – as our main focus for further analysis.

**Table 1.** General assembly statistics and BUSCO scores for RPW genome assemblies.

|  | Assemblies from this study |  |  | Assemblies from Hazzouri *et al.*<sup>13</sup> |  |  |  |
| --- | --- | --- | --- | --- | --- | --- | --- |
|  | 10x pseudo-haplotype1 | 10x pseudo-haplotype2 | 10x diploid megabubbles | Male diploid ABySS | 10x mixed-sex megabubbles | ABYSS+10x (M_v.1) | hybrid (M_pseudochr) |
| Assembly size (bp) | 589,402,552 | 588,821,663 | 730,004,643 | 749,083,425 | 967,735,890 | 780,518,639 | 782,098,041 |
| Assembly size for scaffolds $\geq 250$ bp (bp) | 589,402,552 | 588,821,663 | 730,004,643 | 596,550,086 | 967,735,890 | 780,518,639 | 782,098,041 |
| Contig count (#) | 42,051 | 41,976 | 47,835 | 1,832,276 | 125,191 | 48,516 | 48,516 |
| Contig N50 (bp) | 37,927 | 37,977 | 38,908 | 2,000 | 18,553 | 23,825 | 23,825 |
| Scaffold count (#) | 24,005 | 24,005 | 25,245 | 1,810,484 | 78,408 | 12,400 | 4,807 |
| Scaffold N50 (bp) | 471,583 | 471,583 | 288,817 | 2,289 | 76,013 | 127,745 | 64,117,472 |
| GC content (%) | 32.21 | 32.21 | 32.15 | 32.4 | 32.27 | 32.19 | 32.19 |
| BUSCO (%) |  |  |  |  |  |  |  |
| Complete | 98.1 | 97.5 | 97.3 | 94.9 | 95.3 | 92.8 | 93.2 |
| Single-copy | 96.2 | 95.5 | 71.7 | 93.6 | 13.4 | 16.0 | 54.2 |
| Duplicated | 1.9 | 2.0 | 25.6 | 1.3 | 81.9 | 76.8 | 39.0 |
| Fragmented | 1.1 | 1.1 | 1.1 | 2.9 | 1.6 | 1.4 | 1.3 |
| Missing | 0.8 | 1.4 | 1.6 | 2.2 | 3.1 | 5.8 | 5.5 |

Unphased regions can represent true homozygosity but can also be the result of insufficient linked-read data, and thus differences observed between the pseudo-haplotypes in our assembly possibly underestimate the true heterozygosity in the individual sequenced. To evaluate this possibility, we re-aligned the 10x data generated here to our pseudo-haplotype1 assembly, called single nucleotide variants (SNVs), and visualized the phase blocks together with the B-allele frequency (BAF) of SNVs across the length of each scaffold (Figure 1). This analysis demonstrated that although phased regions largely coincide with blocks of heterozygosity with BAF ~0.5 there are heterozygous regions that remained unphased (e.g. in JAACXV010000004; Figure 1), suggesting that our assembly is underphased most likely because of insufficient linked read data in some regions. Interestingly, this analysis also revealed the presence of large scaffolds with very low density of SNVs (e.g. JAACXV010014584 and JAACXV010014549; Figure 1). Scaffolds lacking heterozygosity could represent homozygous diploid regions resulting from inbreeding or hemizygous sex chromosome sequences if the RPW individual sequenced is male. To investigate these hypotheses and predict the sex of the sequenced individual we mapped female and male Illumina reads from Hazzouri *et al.*^13^ to our pseudo-haplotype1 assembly and calculated the ratio of male/female mean mapped read depth for each scaffold. This analysis allowed us to identify multiple putative sex chromosome scaffolds with a male/female mean mapped read depth ratio of ~0.5 (e.g. JAACXV010000003; Figure 2). We also mapped the DNA-seq reads produced in this study and observed that mean mapped depth of our 10x Genomics reads matches the mapped depth of female reads from Hazzouri *et al.*^13^ in putative sex chromosome scaffolds, indicating that the RPW sample sequenced here is likely to be female and the putative sex chromosomal scaffolds are therefore X-linked. Additionally, the presence of phase blocks and heterozygous regions on these putative X chromosome scaffolds further support the conclusion that these sequences are X-linked and that our sample is female. Using this approach, we identified 29 putative X chromosome scaffolds totaling ~20 Mb (Figure 2). Surprisingly, none of the long scaffolds that are mostly homozygous appear to be sex linked based on their male/female mean mapped read depth ratio (Figure 2, Supplementary File 3), indicating that the RPW genome sequenced here is characterized by long stretches of low heterozygosity that probably arose as a result of inbreeding.

**Figure 1.**
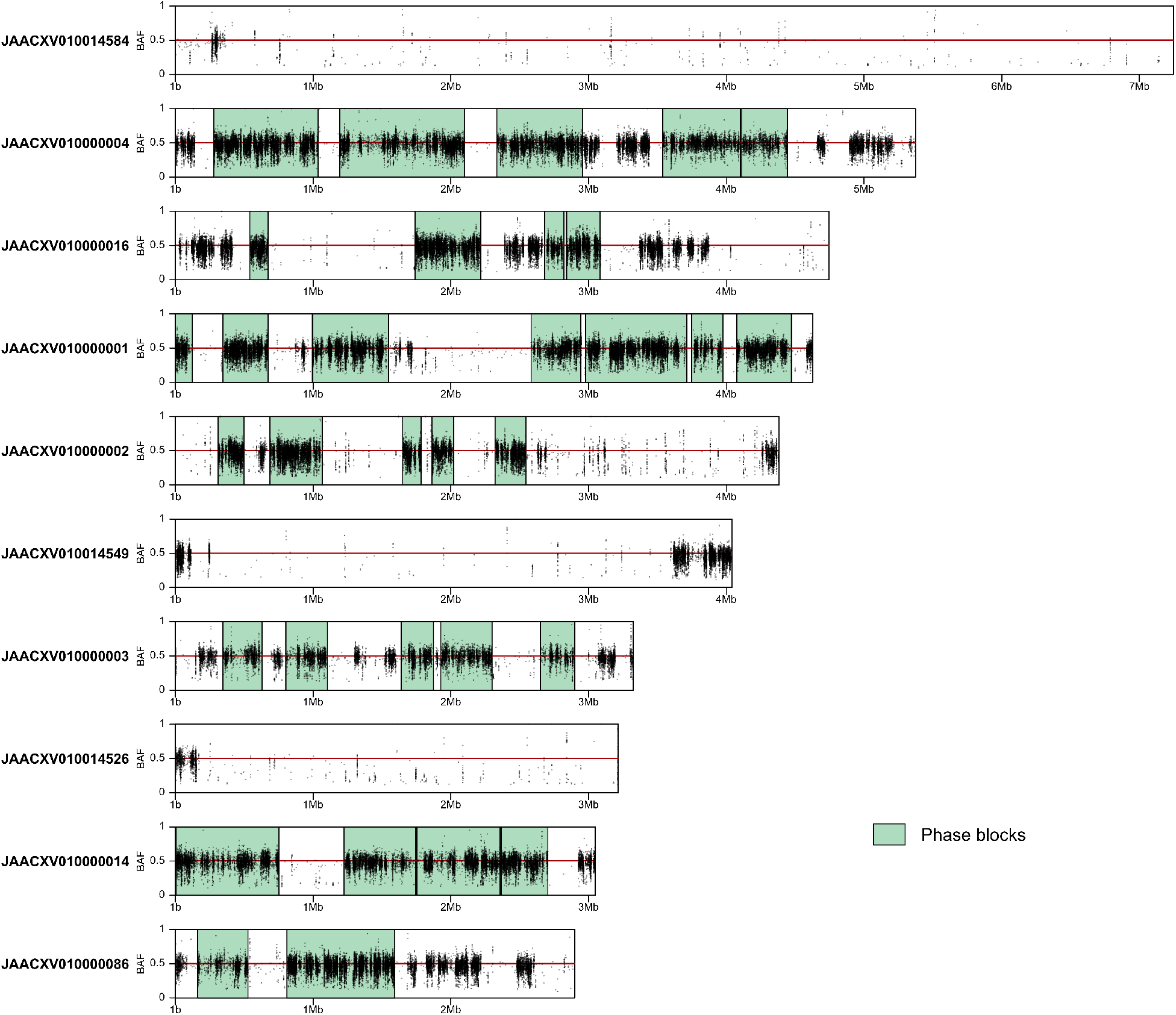
Phase blocks and B-allele frequency (BAF) of single-nucleotide variants (SNVs) in the 10 largest scaffolds of the RPW pseudo-haplotype1 assembly. Phased regions are shown as green highlighted boxes and SNVs as black dots. Regions with white background represent unphased segments of the genome where both pseudo-haplotype assemblies are identical. SNVs in a diploid genome are expected to display BAF values of ~0.5.

**Figure 2.**
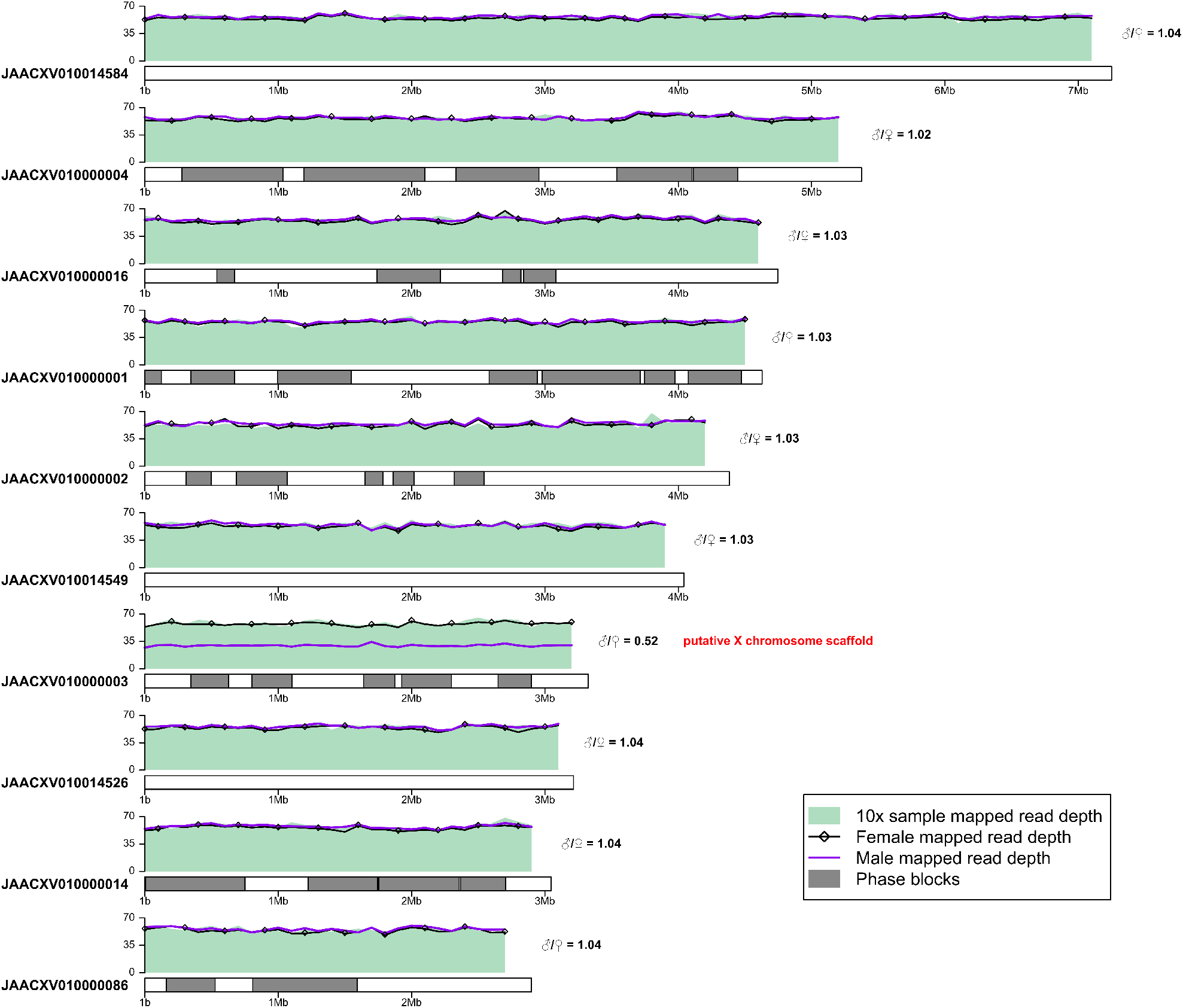
Identification of putative sex chromosome scaffolds. Sequencing data were subsampled to ∼39 Gb and mapped to pseudo-haplotype1. Mean mapped read depth of the 10x Genomics reads produced in this study (SRX7520800; green shaded area), and the female (black line with open diamonds) and male (purple line) Illumina reads from Hazzouri *et al.*^13^ (SRX5416728 and SRX5416729) is shown in 100 kb windows across the 10 longest scaffolds of pseudo-haplotype1 assembly with terminal windows removed. Phase blocks are shown as gray rectangles. The ratio of male/female mean mapped read depth is given on the right side of each scaffold. Scaffolds with a male/female ratio of ∼0.5 are indicated as putative sex chromosome sequences. The similar mapped read depth of our RPW sample and the female sample from Hazzouri *et al.*, as well as the presence of phase blocks on putative sex chromosome scaffolds implies heterozygosity due to diploidy and indicates that such scaffolds are X-linked and that the individual sequenced in this study is female.

Recently, a hybrid assembly constructed using a combination of Illumina, 10x Genomics and Oxford nanopore sequencing has been published by Hazzouri *et al.*^13^. This hybrid assembly was produced by initially scaffolding an ABySS assembly of male Illumina 150 bp PE reads using SSPACE-LongRead^50^ with scaffolds from a Supernova assembly based on a mixed-sex 10x Genomics library (comprised of three males and three females; Khaled Hazzouri, personal communication) exported in ‘megabubbles’ format to create the “M_v.1” hybrid assembly. The resulting M_v.1 hybrid assembly was further merged with two assemblies based on Oxford nanopore data and then scaffolded into pseudochromosomes using synteny with the *T. castaneum* genome to create the “M_pseudochr” assembly available in NCBI (GCA_012979105.1). By using long-read sequences and assuming chromosome-wide synteny and collinearity with *T. castaneum* for scaffolding, the M_pseudochr has a much higher scaffold N50 and also has nearly 200 Mb of additional DNA relative to either of our pseudo-haplotype assemblies (Table 1). However, the contig N50s of both our pseudo-haplotypes are higher than the M_pseudochr assembly. Both pseudo-haplotypes in our assembly also have a higher proportion of complete BUSCOs and a much lower proportion of duplicated BUSCOs than the M_pseudochr assembly (Table 1).

The observation of very high proportion of duplicated genes in the BUSCO gene set for the M_pseudochr assembly is unusual since by definition BUSCO genes are expected to be present in a single copy in most organisms^14^. We hypothesized that the high proportion of duplicated BUSCOs in the M_pseudochr assembly is an artifact of scaffolding the initial male ABySS assembly using a Supernova assembly exported in megabubbles format, which includes multiple haplotypes in a single output file (Supplementary Figure S1). Scaffolding with a megabubbles formatted Supernova assembly could introduce multiple haplotypes of the same locus into a single haploid representation of the genome, thereby creating regions which falsely appear to be duplicated in the M_pseudochr assembly when in fact they are haplotype-induced duplication artifacts^15^. Inclusion of multiple haplotypes in a single haploid assembly could also explain the substantially larger total genome size in the M_pseudochr assembly relative to either of our pseudo-haplotype assemblies (Table 1). Additionally, since the 10x Genomics library used by Hazzouri *et al.*^13^ for Supernova assembly was made from six diploid individuals, heterozygosity will be increased across the genome in this library relative to a single diploid individual.

To provide initial support for the hypothesis that high proportion of duplicated BUSCO genes in the M_pseudochr assembly results from scaffolding with multiple haplotypes from their Supernova megabubbles assembly, we exported our diploid Supernova assembly in megabubbles format and ran BUSCO on the resulting assembly. As predicted, exporting our diploid assembly in megabubbles format led to a much larger total genome size and higher proportion of duplicated BUSCOs (Table 1). We also obtained and analyzed the male ABySS assembly and mixed-sex Supernova megabubbles assemblies used as input to the ABySS+10x (M_v.1) and final hybrid assemblies (M_pseudochr) from Hazzouri *et al.*^13^ (David Nelson, personal communication). As shown in Table 1, their male ABySS assembly has a low proportion of duplicated BUSCO genes, similar to our pseudo-haplotype assemblies. In contrast, their multiple individual mixed-sex Supernova megabubbles assembly has an extremely high proportion of duplicated BUSCO genes, higher even than our diploid Supernova assembly exported in megabubbles format. Their male ABySS assembly has an apparently higher total assembly size than our pseudo-haplotype assemblies, but has a much lower total assembly size (597 Mb) when only scaffolds ≥250 bp are considered (Table 1), suggesting many small scaffolds inflate the total size of their initial male ABySS assembly. Their mixed-sex Supernova megabubbles assembly also has very large total genome size (968 Mb), which is not caused by inclusion of small scaffolds ≤250 bp. A high proportion of duplicated BUSCOs and a large total assembly size are also observed in the M_v.1 hybrid assembly prior to assembling into the final pseudochromosomes (M_pseudochr). Together, these results support the hypothesis that the mixed-sex Supernova megabubbles assembly used for scaffolding by Hazzouri *et al.*^13^ contributed an excess of artifactually-duplicated sequences to their intermediate M_v.1 and final M_pseudochr hybrid assemblies.

Next, we tested which reconstruction of the RPW genome – our pseudo-haplotype1 assembly versus the M_pseudochr hybrid assembly from Hazzouri *et al.*^13^ – has better support in the unassembled DNA-seq data from both projects. To do this, we first classified BUSCO genes as being single copy or duplicated in the M_pseudochr assembly. We then mapped unassembled DNA-seq reads from four datasets (our 10x Genomics library, their 10x Genomics library, their male and female Illumina PE libraries) to our pseudo-haplotype1 assembly. If BUSCO genes classified as duplicated in the M_pseudochr assembly are truly duplicated in the RPW genome but are erroneously collapsed in our pseudo-haplotype1 assembly, we expect these genes to have higher mapped read depth relative to BUSCO genes classified as single-copy. Alternatively, if BUSCO genes classified as duplicated in the M_pseudochr assembly are haplotype-induced duplication artifacts and our pseudo-haplotype assemblies represent the true structure of the RPW genome, we expect no difference in mapped read depth for BUSCO genes classified either as duplicated or single copy in the M_pseudochr assembly. Expectations of the latter hypothesis hold even for the 10x Genomics library from Hazzouri *et al.*^13^ that was generated from multiple individuals if gene copy number is consistent among all individuals in the pooled sample. As shown in Figure 3, despite differences in overall coverage across datasets, we observe no difference in relative mapped read depth for BUSCO genes classified as duplicated versus single copy in the M_pseudochr assembly when DNA-seq reads are mapped to our pseudo-haplotype1 assembly (Kolmogorov-Smirnov Tests; all *P* > 0.05). No difference in read depth for these two categories of BUSCO genes is robustly observed across four different DNA-seq datasets sampled from two geographic locations generated using two different library types, and is not influenced by low quality read mappings (Figure 3). These results indicate that the unassembled DNA-seq data from both projects better support BUSCO gene copy number in our pseudo-haplotype1 reconstruction of the RPW genome.

**Figure 3.**
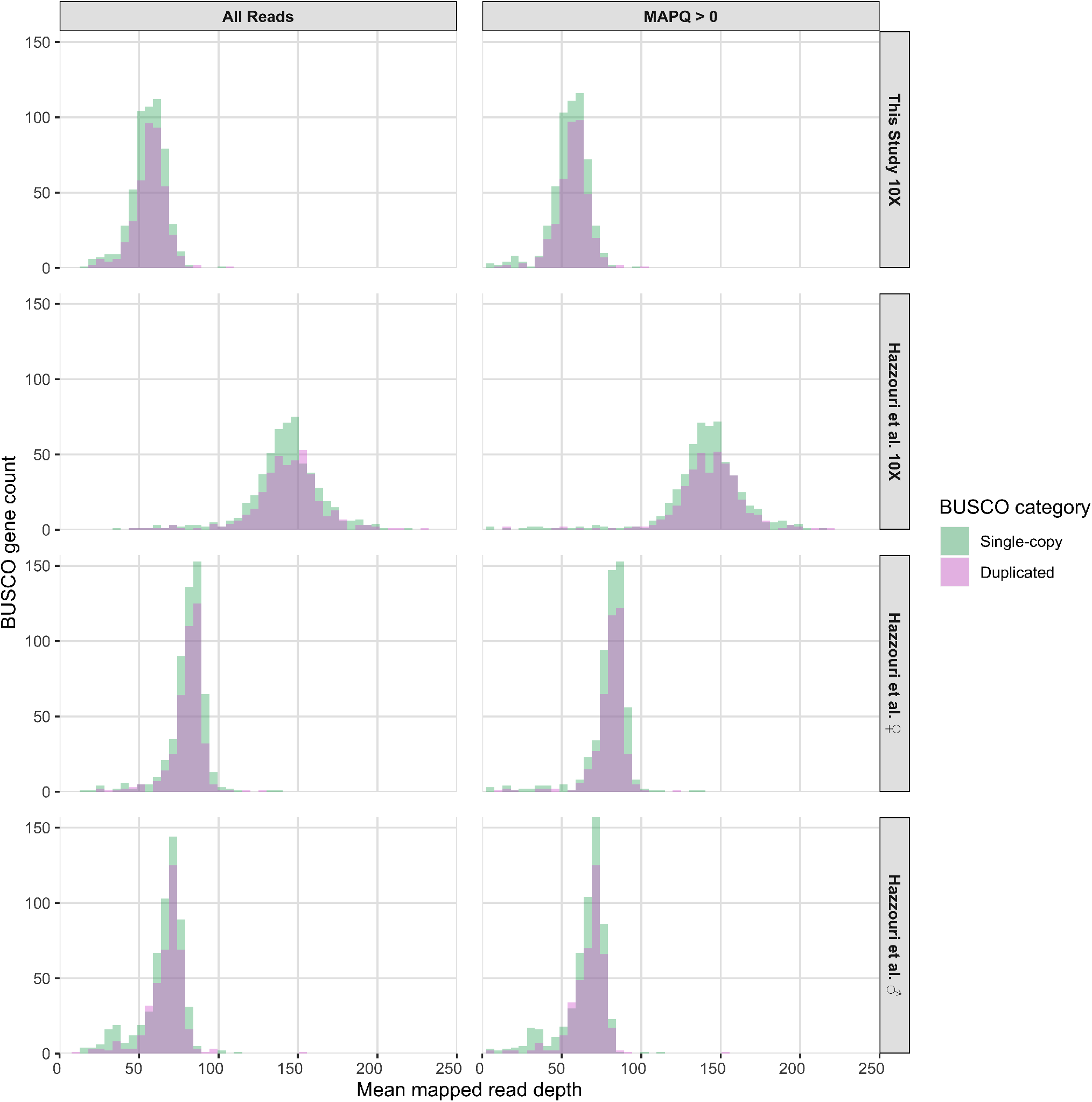
Mapped read depth of BUSCO genes in the RPW pseudo-haplotype1 assembly. BUSCO genes were categorized as single-copy or duplicated based on their status in the M_pseudochr assembly. The four DNA-seq datasets analyzed are (from top to bottom): 10x Genomics library from this study (SRX7520800), 10x Genomics library from Hazzouri *et al.*^13^ (SRX5416727), female RPW Illumina PE library from Hazzouri *et al.*^13^ (SRX5416728), and male RPW Illumina PE library from Hazzouri *et al.*^13^ (SRX5416729). Depth estimates based on all mapped read and high quality mapped reads (MAPQ > 0) are shown in the left and right columns, respectively.

Finally, we estimated total genome size for the RPW using assembly-free k-mer based methods^39, 40^ based on raw DNA-seq reads from our 10x Genomics library and genomic libraries from Hazzouri *et al.*^13^ (Supplementary Table S3; Supplementary Figure S3). Diploid DNA-seq datasets from our study (10x Genomics) and from their male and female Illumina PE libraries all predict a total genome size for the RPW of <600 Mb (Supplementary Table S3), similar to our pseudo-haplotype1 genome assembly. In contrast, their multiple individual mixed-sex 10x Genomics library predicts a much higher genome size than other DNA-seq datasets. However, estimates of genome size based on their multiple individual mixed-sex library are likely biased since is does not fit the assumptions of diploidy required by these methods (Supplementary Figure S3). We note that Hazzouri *et al.*^13^ also reported genome size estimates based on flow cytometry analysis of 726 and 696 Mb for the female and male RPW genome, respectively, which are >100 Mb larger than either of our pseudo-haplotypes assembly sizes (Table 1). This discrepancy could be due to the presence of highly repetitive sequences such as centromeric satellite DNAs and other repeats in the RPW genome that cannot be recovered during genome assembly^51^. Indeed, such differences between physical estimates of genome size and assembly length have been observed in other insects^52, 53^ and often correlate with the abundance of heterochromatic repeats. Integrating all lines of evidence, we conclude that our phased diploid assembly does not include haplotype-induced duplication artifacts and excess sequence observed in the M_v.1 and M_pseudochr hybrid assemblies from Hazzouri *et al.*^13^, and thus provides a more accurate platform for gene discovery in the RPW genome.

### Gene prediction using a haplotype-resolved diploid assembly provides a high-quality RPW gene set

We next performed *ab initio* gene prediction on a repeat-masked version of our pseudo-haplotype1 assembly using BRAKER^19^ trained with protein evidence from multiple insect species and RPW short-read RNA-seq data from three previous studies^7, 9, 10^. Over 45% of pseudo-haplotype1 was identified by RepeatMasker as interspersed repeats using a *de novo* repeat library generated using RepeatModeler and masked prior to annotation. BRAKER annotation of our masked pseudo-haplotype1 assembly predicted 25,382 protein-coding transcripts that clustered into 23,413 loci (Table 2, Supplementary File 4). Over 88% of exon-intron junctions in our BRAKER annotation can be validated by RNA-seq spliced alignments, suggesting that the exon-intron structures for the genes included in our genome annotation have high accuracy. Functional annotation of our BRAKER gene models revealed that ~70% of RPW predicted proteins share similarity to sequences in the NCBI *nr* protein database, and that ~84% had hits to one of InterPro’s member databases. The combined evidence of *nr* and InterPro resulted in over 13,000 (~54%) RPW predicted proteins being annotated with Gene Ontology (GO) terms (Table 3, Supplementary File 5).

**Table 2.** Statistics and BUSCO scores for RPW and *Tribolium* annotation gene sets.

|  | RPW pseudo-haplotype1 BRAKER | RPW isoseq3 | RPW M_v.1 Funannotate | RPW M_pseudochr BRAKER | Tribolium v5.2 |
| --- | --- | --- | --- | --- | --- |
| Transcript (#) | 25,382 | 24,009 | 25,394 | 36,491 | 25,229 |
| Loci (#) | 23,413 | 6,222 | 25,394 | 33,422 | 14,244 |
| BUSCO (%) – all isoforms |  |  |  |  |  |
| Complete | 91.8 | 62.5 | 68.9 | 88.8 | 95.6 |
| Single-copy | 82.0 | 29.8 | 21.1 | 9.6 | 76.2 |
| Duplicated | 9.8 | 32.7 | 47.8 | 79.2 | 19.4 |
| Fragmented | 2.2 | 1.6 | 10.5 | 2.2 | 0.3 |
| Missing | 6.0 | 35.9 | 20.6 | 9.0 | 4.1 |
| BUSCO (%) – one isoform per locus |  |  |  |  |  |
| Complete | 91.2 | 60.1 | 68.9 | 88.3 | 91.6 |
| Single-copy | 89.2 | 59.7 | 21.1 | 10.9 | 91.1 |
| Duplicated | 2.0 | 0.4 | 47.8 | 77.4 | 0.5 |
| Fragmented | 2.2 | 2.6 | 10.5 | 2.2 | 0.5 |
| Missing | 6.6 | 37.3 | 20.6 | 9.5 | 7.9 |

**Table 3.** Functional annotation of 25,382 predicted proteins in the RPW pseudo-haplotype1 annotation.

| Database | Sequences annotated | % of total |
| --- | --- | --- |
| *nr* | 17,893 | 70.5 |
| InterPro | 21,210 | 83.6 |
| *nr* or InterPro | 22,001 | 86.7 |
| Gene Ontology | 13,779 | 54.3 |

BUSCO analysis on the full set of transcripts predicted by BRAKER revealed the presence of 91.8% of Arthropod genes in our annotation, which is close to the upper bound of the total number of Arthropod BUSCOs expected in a well-annotated Coleopteran genome like *T. castaneum* (95.6%) (Table 2). However, our annotation has lower BUSCO completeness relative to the RPW pseudo-haplotype1 sequence itself (98%) (Table 1), which implies that a small proportion of genes present in the assembly are missing from our annotation. The total number of genes in our BRAKER results (23,413 loci) is also substantially higher than the 14,244 loci currently annotated in *T. castaneum*, which may indicate false positive gene models in our BRAKER annotation or real loci in our RPW pseudo-haplotype1 assembly that are split into multiple BRAKER gene models. The total number of loci in our BRAKER annotation is on the same order of the number of RPW loci identified by Hazzouri *et al.*^13^ (25,394), who annotated their intermediate M_v.1 hybrid assembly using Funannotate (https://github.com/nextgenusfs/funannotate). However, when the BRAKER pipeline used to annotate our pseudo-haplotype1 assembly is applied to their final M_pseudochr hybrid assembly, we identifiy a much larger number of loci (33,422) (Table 2).

Both the Funannotate (68.9%) annotation of the M_v.1 assembly performed by Hazzouri *et al.*^13^ and our BRAKER (88.8%) annotation of their M_pseudochr assembly had lower BUSCO completeness than our BRAKER annotation of pseudo-haplotype1 (Table 2). In addition to lower overall BUSCO completeness, both the M_v.1 Funannotate and M_-pseudochr BRAKER annotations have much higher BUSCO duplication than gene sets based on BRAKER annotation of pseudo-haplotype1 or the re-processed Iso-Seq transcriptome (Table 2: “all isoforms”). However, it is important to highlight that the BUSCO method can falsely classify single copy genes as being duplicated when applied to gene sets that include multiple transcript isoforms at the same locus, thereby obscuring the true degree of duplication in a gene set. Therefore, we also performed BUSCO analysis on RPW and *T. castaneum* gene sets using a single isoform selected randomly from each locus (Table 2: “one isoform per locus”). After controlling for the effects of alternative isoforms, 91.2% of Arthropod BUSCOs were captured completely in our BRAKER annotation of pseudo-haplotype1, 89.2% of which were identified as single-copy and only 2% as duplicated. Similarly low rates of duplicated BUSCOs are observed in the RPW Iso-Seq and *T. castaneum* gene sets when the effects of multiple isoforms are eliminated (Table 2). In contrast, even after controlling for the effect of multiple isoforms on estimates of BUSCO gene duplication, we observe very high rates of duplicated BUSCO genes in the M_v.1 Funannotate annotation and the M_pseudochr BRAKER annotation (Table 2). These results indicate that the haplotype-induced duplication artifacts detected in the hybrid genome assemblies from Hazzouri *et al.*^13^ also impact protein-coding gene sets predicted using these genome sequences.

We further evaluated the quality of our BRAKER annotation by comparison to two external datasets of RPW genes. The first dataset is based on a recently-published RPW Iso-Seq transcriptome obtained using PacBio long-read sequences^10^. Preliminary analysis of the processed Iso-Seq dataset reported by Yang *et al.*^10^ mapped to our pseudo-haplotype1 assembly revealed many transcript isoforms on the forward and reverse strands of the same locus (Supplementary Figure S2), presumably due to the inclusion of non-full length cDNA subreads that were sequenced on the anti-sense strand. Therefore, we re-processed CCS reads from Yang *et al.*^10^ using the isoseq3 pipeline and obtained a dataset of 24,136 high-quality transcripts, nearly all of which could be mapped to our pseudo-haplotype1 assembly (24,009, 99.5%). After clustering mapped Iso-Seq transcripts at the genomic level, we identified 6,222 loci supported by this high-quality re-processed Iso-Seq dataset. Overlapping BRAKER annotations with this high-quality Iso-Seq dataset revealed that the vast majority of loci identified using Iso-Seq evidence were also present in our BRAKER annotation (5,875/6,222; 94.4%). This analysis also revealed 6,436 loci in the BRAKER annotation that overlap Iso-Seq loci, with the discrepancy in numbers of overlapping loci among the two datasets explained by one-to-many relationships.

The second external dataset used to evaluate our BRAKER annotation is derived from genes involved in behavior and physiology that may be relevant for RPW pest management^9, 11, 48^. Antony *et al.*^9^ identified 157 putative transcripts belonging to six gene families associated with chemosensation, including: odorant receptors (ORs), odorant binding proteins (OBPs), gustatory receptors (GRs), chemosensory proteins (CSPs), ionotropic receptors (IRs), and sensory neuron membrane proteins (SNMPs). In a later study Antony *et al.*^48^ identified 77 putative transcripts belonging to the cytochrome P450 monooxygenases (CYPs) that are potentially involved in insecticide resistance. Likewise, Zhang *et al.*^11^ identified 88 putative transcripts encoding neuropeptide precursors and their G-protein coupled receptors (GPCRs)^11^. Our pseudo-haplotype1 assembly contains the vast majority of these functionally-relevant RPW genes (306/322, 95%) (Table 4). Only 16 previously reported genes (3 OBPs, 6 CSPs, and 7 CYPs) could not be mapped to our pseudo-haplotype1 assembly. Analysis of unassembled DNA-seq reads revealed that the sequences for these 16 genes are in fact not present in the raw DNA sequencing data produced here or in Hazzouri *et al.*^13^, and thus represent either strain-specific gene variants or assembly/contamination artifacts in the *de novo* transcriptomes from Antony *et al.*^9, 48^. The 306 transcripts that map to our RPW assembly can be clustered into 296 distinct loci (Table 4), indicating that some previously identified transcripts are likely isoforms of the same gene, allelic pairs, or sequencing/assembly artifacts.

**Table 4.** Presence of functionally-relevant gene sets in the RPW pseudo-haplotype1 assembly and annotation.

| Gene family | Transcripts | Transcripts mapped to assembly | Mapped loci | Mapped loci with StringTie models | StringTie models consistent with BRAKER annotation |
| --- | --- | --- | --- | --- | --- |
| OR<sup>†</sup> | 76 | 76 | 74 | 68 | 68 |
| OBP<sup>†</sup> | 38 | 35 | 35 | 34 | 34 |
| GR<sup>†</sup> | 15 | 15 | 15 | 15 | 13 |
| CSP<sup>†</sup> | 12 | 6 | 6 | 6 | 6 |
| IR<sup>†</sup> | 10 | 10 | 10 | 8 | 8 |
| SNMP<sup>†</sup> | 6 | 6 | 6 | 5 | 4 |
| CYP<sup>‡</sup> | 77 | 70 | 64 | 55 | 54 |
| Neuropeptide<sup>§</sup> | 42 | 42 | 40 | 37 | 36 |
| GPCR<sup>§</sup> | 46 | 46 | 46 | 43 | 43 |
| **Total** | 322 | 306 | 296 | 271 | 266 |

<sup>†</sup>Curated gene set from Antony *et al.*<sup>9</sup>. <sup>‡</sup>Curated gene set from Antony *et al.*<sup>48</sup>. <sup>§</sup>Curated gene set from Zhang *et al.*<sup>11</sup>. Numbers of transcripts, numbers of transcripts mapped to the pseudo-haplotype1 assembly, and numbers of loci in the pseudo-haplotype1 assembly are based on the original transcripts reported in Antony *et al.*<sup>9</sup> and Zhang *et al.*<sup>11</sup>. Numbers of loci that are consistent with the BRAKER annotation of pseudo-haplotype1 are based on the corresponding StringTie transcript model overlapping transcripts from Antony *et al.*<sup>9</sup>, Antony *et al.*<sup>48</sup> and Zhang *et al.*<sup>11</sup> to correct for orientation strand orientation artifacts in the Antony *et al.*<sup>9</sup> transcriptome and provide a uniform set of gene structures across curated gene sets.

We initially sought to evaluate the consistency of genomic mappings of transcript models for curated genes against our BRAKER annotation, but preliminary analysis revealed that a majority of chemosensory transcripts in the assembly reported by Antony *et al.*^9^ map to the opposite strand of genes predicted by both BRAKER and our re-processed Iso-Seq transcriptome (Supplementary Figure S2). Opposite strand mappings relative to BRAKER annotations were not observed in the Zhang *et al.*^11^ transcripts or the Antony *et al.*^48^ transcripts. We interpret the opposite strand mapping of many transcripts in the Antony *et al.*^9^ transcriptome assembly as arising from contigs being assembled in the anti-sense orientation. To correct the strand orientation in the Antony *et al.*9 assembly, we generated a reference-guided StringTie assembly of short-read Illumina data and overlapped transcripts from Antony *et al.*^9^ with StringTie transcripts. The majority of curated gene loci overlap StringTie transcripts (271/296; 91.5%) (Table 4). Using the StringTie transcript models as proxies for gene structures that correct strand orientation issues in the Antony *et al.*^9^ assembly and are uniform across curated gene sets, we find that the majority of curated genes are consistent with our BRAKER annotations (266/271; ~98%) (Table 4).

## Conclusion

Here we report a draft diploid assembly and genome-wide gene annotation constructed with the aim of supporting efforts to mitigate the impact of agricultural damage caused by the Red Palm Weevil, *R. ferrugineus*. We demonstrate that phased diploid assembly using 10x Genomics linked reads is a successful strategy for generating scaffolds that are suitable for protein-coding gene prediction in this species. We argue that our haplotype-resolved genome assembly provides a more accurate representation of the *R. ferrugineus* genome that does not include haplotype-induced duplication artifacts present in the genome assembly and annotation reported in a recent study^13^. If our interpretation of the differences between our assembly of the RPW genome and that of Hazzouri *et al.*^13^ is correct, then the claims for high rates of gene family expansion in the RPW lineage reported by Hazzouri *et al.*^13^ could be an artifact of their assembly and should be re-evaluated. Additionally, we show that our annotation of the RPW genome cross-validates the majority of chemosensory and neuropeptide genes previously identified as candidates for guiding management of this pest using molecular genetics^9, 11^, but that a limited number of previously-identified chemosensory genes may be transcriptomic artifacts or strain-specific gene variants. The availability of an RPW genome assembly also allows the identification of strand orientation artifacts in previously-reported transcriptomic datasets for this species^9, 10^. Finally, by integrating our genomic data with rigorously-processed Iso-Seq data^10^, we identify >6,000 RPW loci independently supported by both genome annotation and long-read transcriptomics that represent a high-quality core gene set for future genetic analysis in this economically-important insect pest.

## Data availability

The raw reads used for Supernova genome assembly are available under SRA accession SRX7520800. Pseudo-haplotype1 (primary) and pseudo-haplotype2 (alternate) assemblies are available at GenBank under accession numbers GCA_014462685.1 and GCA_014490705.1, respectively. All other associated data is available in the Supplementary Material and in the Supplementary Files S1-S5 as described in the text.

## Acknowledgments

We thank Noah Workman, Julia A. Portocarrero, and Magdy Alabady at the University of Georgia Genomics and Bioinformatics Core for assistance with 10x Genomics library preparation and Illumina sequencing. We thank Khaled Hazzouri (United Arab Emirates University), David Nelson (New York University Abu Dhabi), and Zhihang Zhou (Hainan University) for providing additional information about published RPW genome resources. We thank Jingxuan Chen, Preston Basting, Shunhua Han, and Claude Desplan for comments on the manuscript. This study was supported in part by resources and technical expertise from the Georgia Advanced Computing Resource Center, and funding from the University of Georgia Research Foundation, and King Abdulaziz City for Science and Technology, Saudi Arabia.

## Author contributions statement

M.M.M., M.B.A-F and C.M.B conceived the experiments, M.A.A., H.A.F.E-S, and F.M.A. conducted the experiments, G.D., C.M.B, and M.M.M. performed the bioinformatic analyses and analyzed the results, G.D., C.M.B, and M.M.M. drafted the manuscript. All authors reviewed the manuscript.

## References

1. Stork, N. E., McBroom, J., Gely, C. & Hamilton, A. J. New approaches narrow global species estimates for beetles, insects, and terrestrial arthropods. Proc Natl Acad Sci USA 112, 7519–7523, DOI: 10.1073/pnas.1502408112 (2015).

2. McKenna, D. D. Beetle genomes in the 21st century: prospects, progress and priorities. Curr. Opin. Insect Sci. 25, 76–82, DOI: 10.1016/j.cois.2017.12.002 (2018).

3. El-Sabea, A. M. R., Faleiro, J. R. & Abo-El-Saad, M. M. The threat of red palm weevil Rhynchophorus ferrugineus to date plantations of the Gulf region in the Middle-East: An economic perspective. Outlooks on Pest Manag. 20, 131–134, DOI: 10.1564/20jun11 (2009).

4. Murphy, S. & Briscoe, B. The red palm weevil as an alien invasive: biology and the prospects for biological control as a component of IPM A Threat to Palms. Biocontrol News Inf 20(1999).

5. Barkan, S., Hoffman, A., Hezroni, A. & Soroker, V. Flight performance and dispersal potential of red palm weevil estimated by repeated flights on flight mill. J Insect Behav 31, 66–82, DOI: 10.1007/s10905-017-9660-y (2018).

6. Wang, L. et al. A large-scale gene discovery for the red palm weevil Rhynchophorus ferrugineus (Coleoptera: Curculionidae). Insect Sci 20, 689–702, DOI: 10.1111/j.1744-7917.2012.01561.x (2013).

7. Yan, W., Liu, L., Qin, W. Q., Li, C. X. & Peng, Z. Q. Transcriptomic identification of chemoreceptor genes in the red palm weevil Rhynchophorus ferrugineus. Genet. Mol Res 14, 7469–7480, DOI: 10.4238/2015.July.3.23 (2015).

8. Yin, A. et al. Transcriptomic study of the red palm weevil Rhynchophorus ferrugineus embryogenesis. Insect Sci 22, 65–82, DOI: 10.1111/1744-7917.12092 (2015).

9. Antony, B. et al. Identification of the genes involved in odorant reception and detection in the palm weevil Rhynchophorus ferrugineus, an important quarantine pest, by antennal transcriptome analysis. BMC Genomics 17, 69, DOI: 10.1186/s12864-016-2362-6 (2016).

10. Yang, H., Xu, D., Zhuo, Z., Hu, J. & Lu, B. SMRT sequencing of the full-length transcriptome of the Rhynchophorus ferrugineus (Coleoptera: Curculionidae). PeerJ 8, e9133, DOI: 10.7717/peerj.9133 (2020).

11. Zhang, H. et al. Neuropeptides and G-protein coupled receptors (GPCRs) in the red palm weevil Rhynchophorus ferrugineus Olivier (Coleoptera: Dryophthoridae). Front Physiol 11, DOI: 10.3389/fphys.2020.00159 (2020).

12. Zhang, X., Wu, R., Wang, Y., Yu, J. & Tang, H. Unzipping haplotypes in diploid and polyploid genomes. Comput. Struct. Biotechnol. J. 18, 66–72, DOI: 10.1016/j.csbj.2019.11.011 (2020).

13. Hazzouri, K. M. et al. The genome of pest Rhynchophorus ferrugineus reveals gene families important at the plant-beetle interface. Commun. Biol. 3, 1–14, DOI: 10.1038/s42003-020-1060-8 (2020).

14. Simao, F. A., Waterhouse, R. M., Ioannidis, P., Kriventseva, E. V. & Zdobnov, E. M. BUSCO: assessing genome assembly and annotation completeness with single-copy orthologs. Bioinformatics 31, 3210–3212, DOI: 10.1093/bioinformatics/btv351 (2015).

15. Kelley, D. R. & Salzberg, S. L. Detection and correction of false segmental duplications caused by genome mis-assembly. Genome Biol 11, R28, DOI: 10.1186/gb-2010-11-3-r28 (2010).

16. Miller, S. A., Dykes, D. D. & Polesky, H. F. A simple salting out procedure for extracting DNA from human nucleated cells. Nucleic Acids Res 16, 1215–1215, DOI: 10.1093/nar/16.3.1215 (1988).

17. Weisenfeld, N. I., Kumar, V., Shah, P., Church, D. M. & Jaffe, D. B. Direct determination of diploid genome sequences. Genome Res 27, 757–767, DOI: 10.1101/gr.214874.116 (2017).

18. Gremme, G., Steinbiss, S. & Kurtz, S. GenomeTools: a comprehensive software library for efficient processing of structured genome annotations. IEEE/ACM Transactions on Comput. Biol. Bioinforma. 10, 645–656, DOI: 10.1109/TCBB.2013.68 (2013).

19. Brůna, T., Hoff, K. J., Lomsadze, A., Stanke, M. & Borodovsky, M. BRAKER2: Automatic eukaryotic genome annotation with GeneMark-EP+ and AUGUSTUS supported by a protein database. bioRxiv 2020.08.10.245134, DOI: 10.1101/2020.08.10.245134 (2020).

20. Stanke, M. et al. AUGUSTUS: Ab initio prediction of alternative transcripts. Nucleic Acids Res 34, W435–439, DOI: 10.1093/nar/gkl200 (2006).

21. Stanke, M., Schoffmann, O., Morgenstern, B. & Waack, S. Gene prediction in eukaryotes with a generalized hidden Markov model that uses hints from external sources. BMC Bioinforma. 7, 62, DOI: 10.1186/1471-2105-7-62 (2006).

22. Stanke, M., Diekhans, M., Baertsch, R. & Haussler, D. Using native and syntenically mapped cDNA alignments to improve de novo gene finding. Bioinformatics 24, 637–644, DOI: 10.1093/bioinformatics/btn013 (2008).

23. Chen, S., Zhou, Y., Chen, Y. & Gu, J. fastp: an ultra-fast all-in-one FASTQ preprocessor. Bioinformatics 34, i884–i890, DOI: 10.1093/bioinformatics/bty560 (2018).

24. Kim, D., Paggi, J. M., Park, C., Bennett, C. & Salzberg, S. L. Graph-based genome alignment and genotyping with HISAT2 and HISAT-genotype. Nat Biotechnol 37, 907–915, DOI: 10.1038/s41587-019-0201-4 (2019).

25. Li, H. et al. The Sequence Alignment/Map format and SAMtools. Bioinformatics 25, 2078–2079, DOI: 10.1093/bioinformatics/btp352 (2009).

26. Bruna, T., Lomsadze, A. & Borodovsky, M. GeneMark-EP+: Eukaryotic gene prediction with self-training in the space of genes and proteins. NAR Genom Bioinform 2, DOI: 10.1093/nargab/lqaa026 (2020).

27. Jones, P. et al. InterProScan 5: genome-scale protein function classification. Bioinformatics 30, 1236–1240, DOI: 10.1093/bioinformatics/btu031 (2014).

28. Buchfink, B., Xie, C. & Huson, D. H. Fast and sensitive protein alignment using DIAMOND. Nat Methods 12, 59–60, DOI: 10.1038/nmeth.3176 (2015).

29. Gotz, S. et al. High-throughput functional annotation and data mining with the Blast2GO suite. Nucleic Acids Res 36, 3420–3435, DOI: 10.1093/nar/gkn176 (2008).

30. Ashburner, M. et al. Gene Ontology: tool for the unification of biology. Nat. Genet. 25, 25–29, DOI: 10.1038/75556 (2000).

31. Shelton, J. M. et al. Tools and pipelines for BioNano data: molecule assembly pipeline and FASTA super scaffolding tool. BMC Genomics 16, 734, DOI: 10.1186/s12864-015-1911-8 (2015).

32. Bushnell, B. BBMap: a fast, accurate, splice-aware aligner. Tech. Rep. LBNL-7065E, Lawrence Berkeley National Lab. (LBNL), Berkeley, CA (United States) (2014).

33. Kriventseva, E. V. et al. OrthoDB v10: sampling the diversity of animal, plant, fungal, protist, bacterial and viral genomes for evolutionary and functional annotations of orthologs. Nucleic Acids Res 47, D807–D811, DOI: 10.1093/nar/gky1053 (2019).

34. Li, H. Minimap2: pairwise alignment for nucleotide sequences. Bioinformatics 34, 3094–3100, DOI: 10.1093/bioinformatics/bty191 (2018).

35. Danecek, P. et al. The variant call format and VCFtools. Bioinformatics 27, 2156–2158, DOI: 10.1093/bioinformatics/btr330 (2011).

36. Gel, B. & Serra, E. karyoploteR: an R/Bioconductor package to plot customizable genomes displaying arbitrary data. Bioinformatics 33, 3088–3090, DOI: 10.1093/bioinformatics/btx346 (2017).

37. Li, H. Aligning sequence reads, clone sequences and assembly contigs with BWA-MEM. arXiv 1303.3997 (2013).

38. Quinlan, A. R. & Hall, I. M. BEDTools: a flexible suite of utilities for comparing genomic features. Bioinformatics 26, 841–842, DOI: 10.1093/bioinformatics/btq033 (2010).

39. Sun, H., Ding, J., Piednoël, M. & Schneeberger, K. findGSE: estimating genome size variation within human and Arabidopsis using k-mer frequencies. Bioinformatics 34, 550–557, DOI: 10.1093/bioinformatics/btx637 (2018).

40. Vurture, G. W. et al. GenomeScope: fast reference-free genome profiling from short reads. Bioinformatics 33, 2202–2204, DOI: 10.1093/bioinformatics/btx153 (2017).

41. Marcais, G. & Kingsford, C. A fast, lock-free approach for efficient parallel counting of occurrences of k-mers. Bioinformatics 27, 764–770, DOI: 10.1093/bioinformatics/btr011 (2011).

42. R Core Team. R: A Language and Environment for Statistical Computing (R Foundation for Statistical Computing, Vienna, Austria, 2017).

43. Wickham, H. ggplot2: Elegant Graphics for Data Analysis (Springer-Verlag New York, 2016).

44. Pertea, G. & Pertea, M. GFF Utilities: GffRead and GffCompare. F1000Res 9, 304, DOI: 10.12688/f1000research.23297.1 (2020).

45. Salmela, L. & Rivals, E. LoRDEC: accurate and efficient long read error correction. Bioinformatics 30, 3506–3514, DOI: 10.1093/bioinformatics/btu538 (2014).

46. Hu, R., Sun, G. & Sun, X. LSCplus: a fast solution for improving long read accuracy by short read alignment. BMC Bioinforma. 17, 451, DOI: 10.1186/s12859-016-1316-y (2016).

47. Kuhn, R. M., Haussler, D. & Kent, W. J. The UCSC genome browser and associated tools. Brief Bioinform 14, 144–161, DOI: 10.1093/bib/bbs038 (2013).

48. Antony, B. et al. Global transcriptome profiling and functional analysis reveal that tissue-specific constitutive overexpression of cytochrome P450s confers tolerance to imidacloprid in palm weevils in date palm fields. BMC Genomics 20, 440, DOI: 10.1186/s12864-019-5837-4 (2019).

49. Pertea, M. et al. StringTie enables improved reconstruction of a transcriptome from RNA-seq reads. Nat. Biotechnol. 33, 290–295, DOI: 10.1038/nbt.3122 (2015).

50. Boetzer, M. & Pirovano, W. SSPACE-LongRead: scaffolding bacterial draft genomes using long read sequence information. BMC Bioinforma. 15, 211, DOI: 10.1186/1471-2105-15-211 (2014).

51. Treangen, T. J. & Salzberg, S. L. Repetitive DNA and next-generation sequencing: computational challenges and solutions. Nat. Rev. Genet. 13, 36–46, DOI: 10.1038/nrg3117 (2012).

52. Bosco, G., Campbell, P., Leiva-Neto, J. T. & Markow, T. A. Analysis of Drosophila species genome size and satellite DNA content reveals significant differences among strains as well as between species. Genetics 177, 1277–1290, DOI: 10.1534/genetics.107.075069 (2007).

53. Pflug, J. M., Holmes, V. R., Burrus, C., Johnston, J. S. & Maddison, D. R. Measuring genome sizes using read-depth, k-mers, and flow cytometry: methodological comparisons in beetles (Coleoptera). G3 10, 3047–3060, DOI: 10.1534/g3.120.401028 (2020).

