## Supplemental Tables and Figures for "Haplotype-resolved genome assembly enables gene discovery in the red palm weevil *Rhynchophorus ferrugineus*"

Supplementary Table S1: **Species and genome assemblies used for protein sequence evidence in the BRAKER annotation of RPW pseudo-haplotype1.**

| Species | Order | Family | Accession |
|---|---|---|---|
| *Aethina tumida* | Coleoptera | Nitidulidae | GCF_001937115.1 |
| *Agrilus planipennis* | Coleoptera | Buprestidae | GCF_000390285.2 |
| *Anoplophora glabripennis* | Coleoptera | Cerambycidae | GCF_000699045.2 |
| *Asbolus verrucosus* | Coleoptera | Tenebrionidae | GCA_004193795.1 |
| *Callosobruchus maculatus* | Coleoptera | Chrysomelidae | GCA_900659725.1 |
| *Drosophila melanogaster* | Diptera | Drosophilidae | GCF_000001215.4 |
| *Dendroctonus ponderosae* (female) | Coleoptera | Curculionidae | GCA_000346045.2 |
| *Dendroctonus ponderosae* (male) | Coleoptera | Curculionidae | GCF_000355655.1 |
| *Diabrotica virgifera* | Coleoptera | Chrysomelidae | GCF_003013835.1 |
| *Ignelater luminosus* | Coleoptera | Elateridae | GCA_011009095.1 |
| *Leptinotarsa decemlineata* | Coleoptera | Chrysomelidae | GCF_000500325.1 |
| *Nicrophorus vespilloides* | Coleoptera | Silphidae | GCF_001412225.1 |
| *Onthophagus taurus* | Coleoptera | Scarabaeidae | GCF_000648695.1 |
| *Oryctes borbonicus* | Coleoptera | Scarabaeidae | GCA_001443705.1 |
| *Photinus pyralis* | Coleoptera | Lampyridae | GCF_008802855.1 |
| *Sitophilus oryzae* | Coleoptera | Curculionidae | GCF_002938485.1 |
| *Tribolium castaneum* | Coleoptera | Tenebrionidae | GCF_000002335.3 |

Supplementary Table S2: **Summary of nucleotide differences between RPW pseudo-haplotype1 and pseudo-haplotype2 assemblies.**

| | ≥1 kb | | ≥10 kb | | ≥50 kb | |
|---|---|---|---|---|---|---|
| Variant type | # differences | differences / kb | # differences | differences / kb | # differences | differences / kb |
| Substitution | 2,056,517 | 3.5837 | 1,950,309 | 3.9107 | 1,078,482 | 3.0011 |
| 1 bp deletion | 48,050 | 0.0837 | 45,656 | 0.0915 | 25,720 | 0.0716 |
| 1 bp insertion | 47,731 | 0.0832 | 45,382 | 0.0910 | 25,550 | 0.0711 |
| 2 bp deletion | 16,522 | 0.0288 | 15,751 | 0.0316 | 8,917 | 0.0248 |
| 2 bp insertion | 16,286 | 0.0284 | 15,520 | 0.0311 | 8,776 | 0.0244 |
| 3-50 bp deletion | 61,144 | 0.1066 | 57,941 | 0.1162 | 32,540 | 0.0906 |
| 3-50 bp insertion | 61,557 | 0.1073 | 58,399 | 0.1171 | 32,694 | 0.0910 |
| 50-1000 bp deletion | 9,864 | 0.0172 | 9,363 | 0.0188 | 5,233 | 0.0146 |
| 50-1000 bp insertion | 9,519 | 0.0166 | 9,057 | 0.0182 | 5,007 | 0.0139 |
| 1 kb+ insertion | 375 | 0.0007 | 370 | 0.0007 | 211 | 0.0006 |
| 1 kb+ deletion | 358 | 0.0006 | 347 | 0.0007 | 195 | 0.0005 |

Numbers of absolute differences and numbers of differences per 1 kb are shown for orthologous minimap2 alignments longer than 1 kb, 10 kb and 50 kb, respectively. Inclusion of shorter alignments increases the absolute number of differences identified, but the corresponding increase in genomic regions covered leads to only minor changes in normalized levels of differences between pseudo-haplotypes. The total aligned sequence length is 573,852,952 bp for alignments ≥1 kb, 498,705,929 bp for alignments ≥10 kb, and 359,359,388 bp for alignments ≥50 kb, respectively.

Supplementary Table S3: **K-mer based genome size estimates (in bp) from unassembled RPW DNA-seq data.**

| Method | This study 10x | Hazzouri et al.[1] 10x | Hazzouri *et al.*[1] ♀ | Hazzouri *et al.*[1] ♂ |
|---|---|---|---|---|
| findGSE | 590,055,225 | 929,305,983† | 592,913,934 | 569,174,629 |
| GenomeScope | 573,088,616 | 1,420,637,199† | 577,505,135 | 554,924,223 |

† Genome size estimates based on the 10x Genomics library from Hazzouri *et al.*[1] (SRX5416727) are biased since this library was generated from multiple individuals and thus it does not fit the assumptions of k-mer based genome size estimation methods that assume diploidy.[2,3]

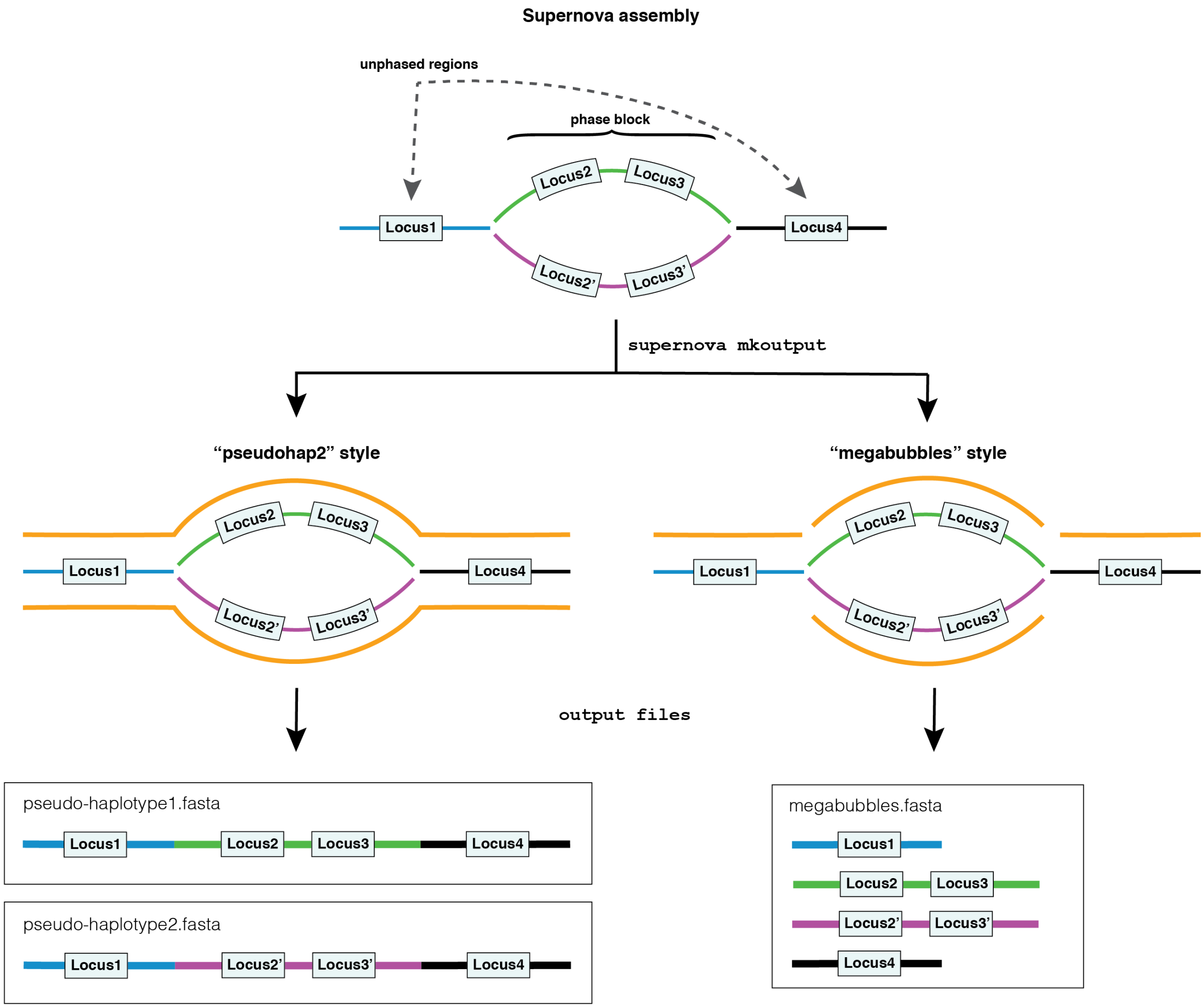


Supplementary Figure S1: **Overview of Supernova assembly output formats.** Supernova assemblies of 10x Genomics libraries can be exported in different output formats that represent phased and unphased regions of the genome in different styles.[4] The "pseudohap2" style generates two distinct "pseudo-haplotype" assembly files that differ in genomic regions where maternal and paternal haplotypes can be phased ("phase blocks") but are identical in homozygous and unphased blocks of the genome. Phasing occurs independently in each phase block and information about the linkage of maternal and paternal phase blocks does not span unphased regions. Thus, the resulting haploid assemblies contain a mixture of maternal and paternal segments and are considered pseudo-haplotypes rather than true haplotypes. Importantly, only one copy of each locus is included per pseudo-haplotype output file. The "megabubbles" style generates one assembly that includes maternal and paternal phase blocks together with unphased blocks in a single file. A megabubbles formatted Supernova assembly thus contains multiple haplotypes for the same locus in a single file, which can lead to haplotype-induced duplication artifacts when this format is used for scaffolding or analyzing gene content.

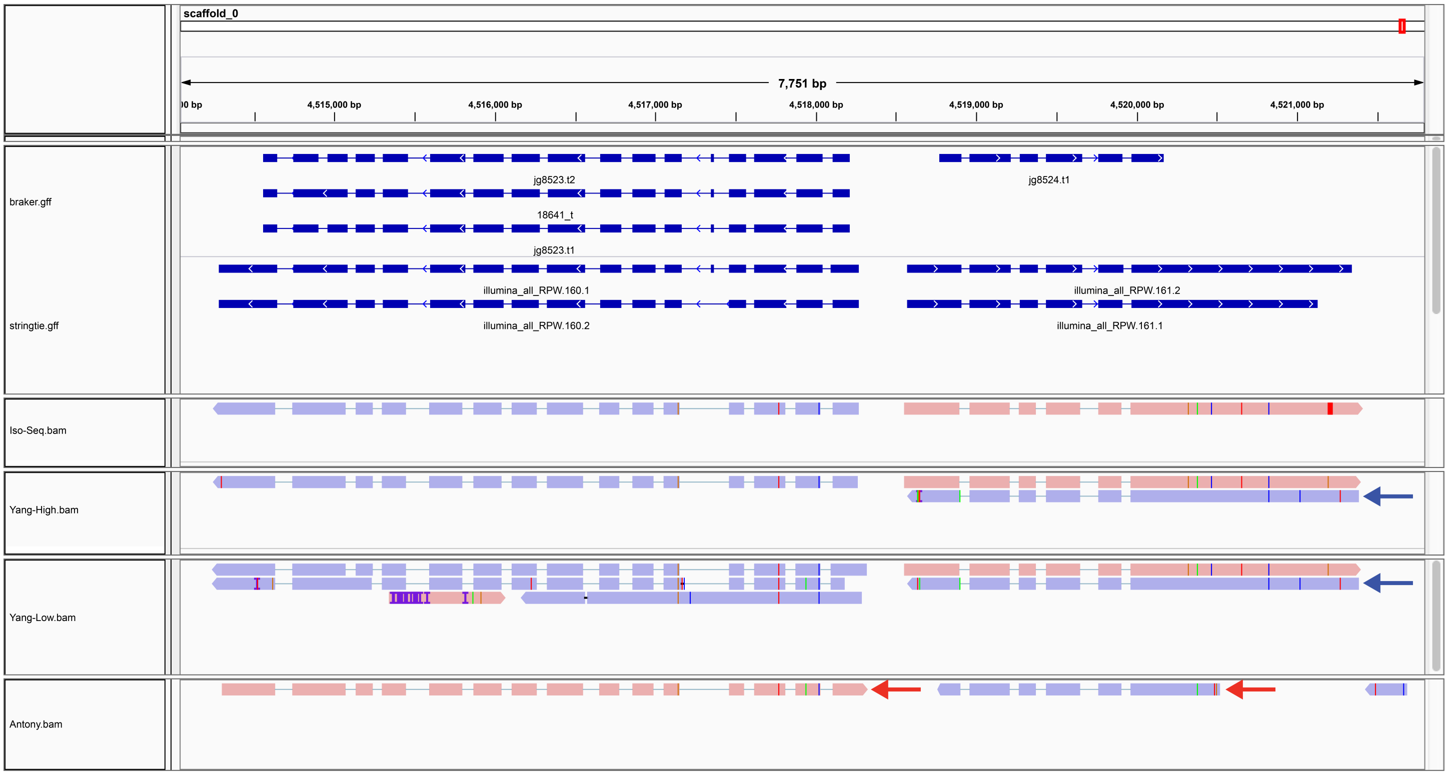


Supplementary Figure S2: **Integrated Genomics Viewer screenshot of strand orientation artifacts in published RPW transcriptomes.** Tracks from top to bottom are: transcript models from the BRAKER annotation of pseudo-haplotype1; StringTie transcript models based on short read RNA-seq; transcripts from our re-processing of long-read Iso-Seq data from Yang *et al.*;[5] transcripts from our the "high-quality" long-read Iso-Seq dataset from Yang *et al.*[5] (SRX7519788); transcripts from the "low-quality" long-read Iso-Seq dataset from Yang *et al.*[5] (SRX8694670); transcripts from the short-read transcriptome assembly from Antony *et al.*[6] (GDKA00000000). Transcript orientation is represented by arrowheads in the top two tracks and by color (red: forward; blue: reverse) and terminal exon direction in the bottom four tracks. Dark blue arrows highlight transcripts in the processed datasets from Yang *et al.*[5] that map to the opposite strand as BRAKER and StringTie models, and other Iso-Seq transcripts from the same locus. We interpret these to represent non-full length cDNA subreads that were sequenced on the anti-sense strand that were not filtered out in the Yang *et al.*[5] processed transcriptomes, but which are properly filtered in our isoseq3 re-processed version of the data from Yang *et al.*[5] Dark red arrows highlight transcripts in the transcriptome assembly from Antony *et al.*[6] that were assembled in the anti-sense orientation relative to BRAKER and StringTie models.

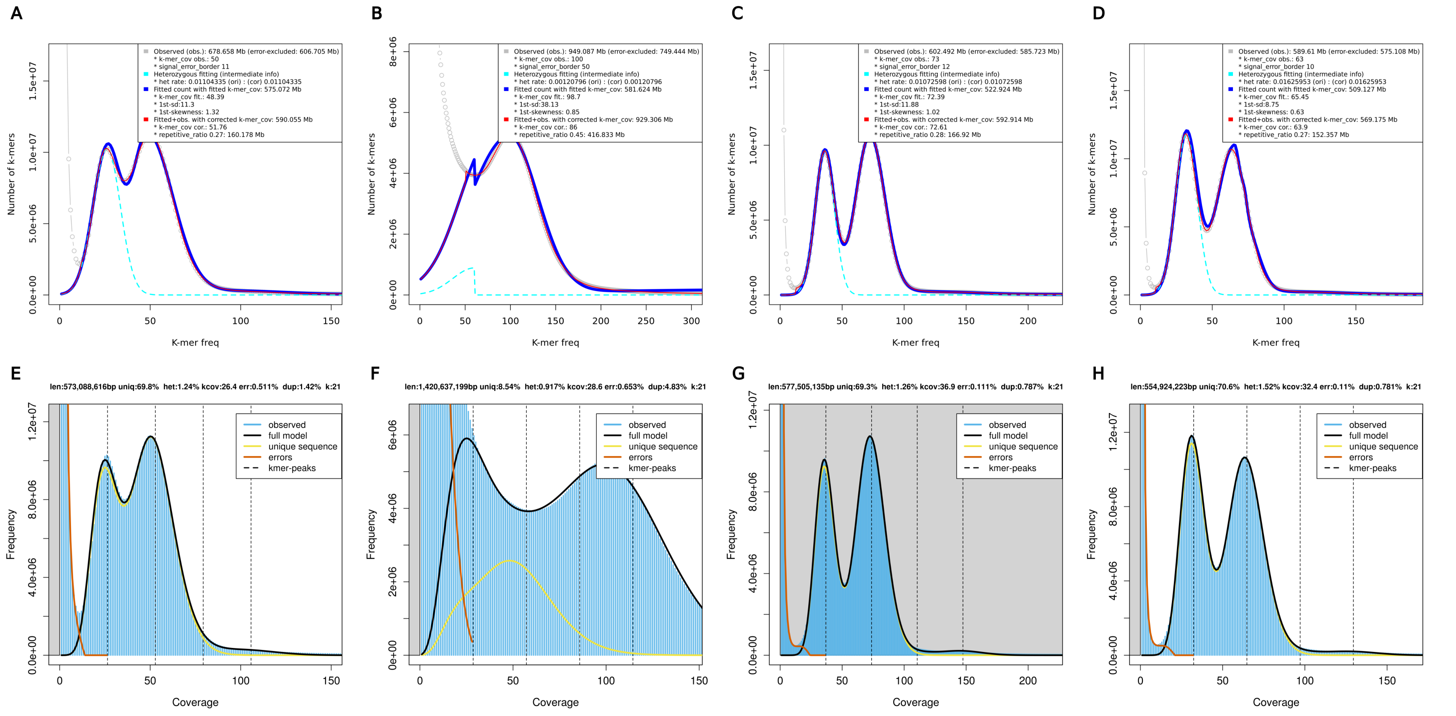


Supplementary Figure S3: **K-mer frequency profiles and genome size estimates for the RPW using unassembled DNA-seq reads.** Rows correspond to the two methods used for genome size estimation: (A-D) findGSE,[2] and (E-H) GenomeScope.[3] Columns correspond to the four RPW DNA-seq datasets analyzed and are arranged as follows: (A, E) 10x Genomics library from this study (SRX7520800); (B, F) 10x Genomics library from Hazzouri *et al.*[1] (SRX5416727); (C, G) female Illumina PE library from Hazzouri *et al.*[1] (SRX5416728); and (D, H) male Illumina PE library from Hazzouri *et al.*[1] (SRX5416729). All estimates were obtained from 21-mer frequency histograms with a max depth of 1,000,000. We note that estimates based on the 10x Genomics library from Hazzouri *et al.*[1] (SRX5416727) are biased since it was generated from multiple individuals and thus does not fit the assumptions of k-mer based genome size estimation methods that assume diploidy.[2,3]

### References

[1] Khaled Michel Hazzouri, Naganeeswaran Sudalaimuthuasari, Biduth Kundu, David Nelson, Mohammad Ali Al-Deeb, Alain Le Mansour, Johnston J. Spencer, Claude Desplan, and Khaled M. A. Amiri. The genome of pest Rhynchophorus ferrugineus reveals gene families important at the plant-beetle interface. *Communications Biology*, 3(1):1–14, June 2020.

[2] Hequan Sun, Jia Ding, Mathieu Piednoël, and Korbinian Schneeberger. findGSE: estimating genome size variation within human and Arabidopsis using k-mer frequencies. *Bioinformatics*, 34(4):550–557, 2018.

[3] Gregory W. Vurture, Fritz J. Sedlazeck, Maria Nattestad, Charles J. Underwood, Han Fang, James Gurtowski, and Michael C. Schatz. GenomeScope: fast reference-free genome profiling from short reads. *Bioinformatics*, 33(14):2202–2204, July 2017.

[4] Neil I. Weisenfeld, Vijay Kumar, Preyas Shah, Deanna M. Church, and David B. Jaffe. Direct determination of diploid genome sequences. *Genome Res*, 27(5):757–767, 2017.

[5] Hongjun Yang, Danping Xu, Zhihang Zhuo, Jiameng Hu, and Baoqian Lu. SMRT sequencing of the full-length transcriptome of the Rhynchophorus ferrugineus (Coleoptera: Curculionidae). *PeerJ*, 8:e9133, May 2020.

[6] Binu Antony, Alan Soffan, Jernej Jakše, Mahmoud M. Abdelazim, Saleh A. Aldosari, Abdulrahman S. Aldawood, and Arnab Pain. Identification of the genes involved in odorant reception and detection in the palm weevil Rhynchophorus ferrugineus, an important quarantine pest, by antennal transcriptome analysis. *BMC Genomics*, 17(1):69, January 2016.
